# Genetic interaction of mammalian IFT-A paralogs regulates cilia disassembly, ciliary protein trafficking, Hedgehog signaling and embryogenesis

**DOI:** 10.1101/803544

**Authors:** Wei Wang, Bailey A. Allard, Tana S. Pottorf, Jay L. Vivian, Pamela V. Tran

## Abstract

Primary cilia are sensory organelles that are essential for eukaryotic development and health. These antenna-like structures are synthesized by intraflagellar transport protein complexes, IFT-B and IFT-A, which mediate bi-directional protein trafficking along the ciliary axoneme. Here using mouse embryonic fibroblasts (MEF), we investigate the ciliary roles of two mammalian orthologues of *Chlamydomonas* IFT-A gene, *IFT139*, namely *Thm1* (also known as *Ttc21b*) and *Thm2 (Ttc21a). Thm1* loss causes perinatal lethality, and *Thm2* loss allows survival into adulthood. At E14.5, the number of *Thm1;Thm2* double mutant embryos is lower than that for a Mendelian ratio, indicating deletion of *Thm1* and *Thm2* causes mid-gestational lethality. We examined the ciliary phenotypes of mutant MEF. *Thm1*-mutant MEF show decreased cilia assembly, shortened primary cilia, a retrograde IFT defect for IFT and BBS proteins, and reduced ciliary entry of membrane-associated proteins. *Thm1*-mutant cilia also show a retrograde transport defect for the Hedgehog transducer, Smoothened, and an impaired response to Smoothened agonist, SAG. *Thm2*-null MEF show normal ciliary dynamics and Hedgehog signaling, but additional loss of a *Thm1* allele impairs response to SAG. Further, *Thm1;Thm2* double mutant MEF show enhanced cilia disassembly, and relative to *Thm1*-null MEF, increased impairment of IFT81 retrograde transport and of INPP5E ciliary import. Thus, *Thm1* and *Thm2* have unique and redundant roles in MEF. *Thm1* regulates cilia assembly, and together with *Thm2*, cilia disassembly. Moreover, *Thm1* alone and together with *Thm2*, regulates ciliary protein trafficking, Hedgehog signaling, and embryogenesis. These findings shed light on mechanisms underlying *Thm1*-, *Thm2*- or IFT-A-mediated ciliopathies.

## Introduction

Cilia are evolutionarily-conserved organelles present in most eukaryotic organisms from *Chlamydomonas reinhardtii* to vertebrates (1). These microtubular organelles can confer motility or act as sensory organelles. In the latter role, a singular cilium, termed a primary cilium, extends from the apical surface of a cell, where it detects and transduces extracellular signals. Primary cilia regulate cell cycle, cell differentiation and cell-cell communication (2).

Mutations that disrupt cilia function cause multi-system disorders, termed ciliopathies (3). Clinical manifestations can include craniofacial and neural tube defects, retinal degeneration, skeletal dysplasia, fibrocystic diseases of the kidney, liver, and pancreas, obesity, and male infertility (4). Understanding the role of ciliary proteins in mediating ciliary processes can provide insight into cellular mechanisms underlying these various defects.

The primary cilium consists of a microtubule-based ciliary axoneme, ensheathed in a ciliary membrane. Extension and maintenance of the axoneme is dependent on multiple protein complexes (5). The IFT-B complex, comprised of approximately 15 proteins, and the kinesin motor mediate anterograde transport, moving protein cargo from the base to the tip of the primary cilium. The IFT-A complex, comprised of approximately 7 proteins, together with the dynein motor mediate retrograde transport, returning cargo from the tip to the base of the primary cilium. IFT-A proteins also mediate ciliary entry of signaling and membrane-associated proteins (6–8). Another protein complex, the BBSome, traffics signaling molecules to the cilium and throughout the cilium where it acts as an adaptor between the IFT complexes and the protein cargoes(9). Despite that these ciliary proteins assemble into complexes, mutations in ciliary genes that encode proteins of the same ciliary complex can result in different phenotypes (10). Thus, individual ciliary proteins likely have cell-specific roles.

Previously, we identified the mammalian IFT-A gene, *Thm1* (TPR-containing Hedgehog Modulator 1; also known as *Ttc21b*), an orthologue of *Chlamydomonas reinhardtii FLA17*/*IFT139* (11, 12). In mouse, early embryonic loss of *Thm1* misregulates Hedgehog (Hh) signaling and causes perinatal lethality, polydactyly, and defects of the skeleton, forebrain, palate, and neural tube (11, 13). Additionally, in mice, *Thm1* deletion in the perinatal period causes renal cystic disease (14), and its deletion in adulthood causes obesity (15). These phenotypes recapitulate the clinical features present in individuals with ciliopathies who have *THM1* mutations. Causative mutations in *THM1* have been identified in patients with nephronophthisis, a renal fibrocystic disease, while modifying mutations in *THM1* have been found in patients with Bardet Biedl Syndrome, which manifests obesity as a cardinal clinical feature (16).

Loss or deficiency of *Thm1* impairs retrograde IFT, resulting in accumulation of IFT proteins and signaling molecules at the ciliary distal tip (11, 17). In some cells, such as mesenchymal cells of the developing limb bud, *Thm1* depletion causes shortened cilia (11), while in cultured RPE cells, cilia length was not affected (17). In *Chlamydomonas Ift139-null* mutants, levels of other IFT-A proteins were reduced, indicating that in the green alga, IFT139 is required for assembly of the IFT-A complex (18).

While there is only one *IFT139* gene in *Chlamydomonas*, evolution has generated two vertebrate orthologues, which we have called *Thm1* and *Thm2*. THM1 and THM2 are both 50% identical to *Chlamydomonas* IFT139 (11, 12). Additionally, the murine *Thm1* and *Thm2* sequences are 50% homologous and encode proteins that have very similar predicted protein structures with multiple (10–11) tetratricopeptide repeat (TPR) domains. At embryonic day (E) 10.5, *Thm1* and *Thm2* also show very similar RNA expression patterns in whole mouse embryos. Recently, *THM2* mutations were reported in adult males with subfertility (19). Still the role of *Thm2* in ciliogenesis and development remains uncharacterized. By generating a *Thm2*-null mouse and using *Thm1*- and *Thm2*-null derived mouse embryonic fibroblasts (MEF), here we show a role for *Thm1* in cilia assembly, and an interaction between *Thm1* and *Thm2* that is important for cilia disassembly, ciliary protein trafficking, Hh signaling and embryogenesis.

## Materials and Methods

### Generation of Thm2 knockout (Thm2^−/−^) mouse

*Thm2*-null mice were generated using CRISPR/Cas9 genome editing. Two guide RNAs (gRNAs) – one targeting exon 4 (target sequence CATACTCCCTGGCCTTGTCGTGG) and the other in intron 8 (target sequence AACCTGACCGACAGCCCACCTGG) - were designed to delete exons 4-8 and ultimately cause a premature stop codon. gRNAs were generated and validated by the Washington University Genome Engineering and iPSC Center). Pro-nuclear injections of 20ng/μl of each in vitro transcribed gRNA and 50ng/μl of Cas9 mRNA of 317 F1 embryos derived from FVB oocyte donors and C57BL6/J males yielded 42 founders. Genomic DNA of founders was amplified and sequenced to determine presence of exon 4-8 deletion and a resulting premature stop codon. Founders carrying a large deletion and stop codon were then crossed to FVB mice to expand the *Thm2−/−* lines.

### Genotyping of NHEJ events following CRISPR/Cas9 genome editing

Three PCR primer sets were designed to characterize the deletions generated by non-homologous end joining (NHEJ) following CRISPR/Cas9 genome editing. Two sets of primers were designed around the exon 4 gRNA, and one set of primers was designed to flank the entire region between exon 4 and intron 8. The first primer set surrounding exon 4 included F-Thm2ex4p (5’-TAC TAC GCC AGC CTC TTC CT-3’) and R-Thm2ex4p (5’-CCC TCC TGT ACC TCT TTG GA-3’) and was expected to produce a WT band of 107bp. The second primer set surrounding exon 4 included F-Thm2ex4r (5’-TGT CTG AAG CCA ACA GAG AGG-3’) and R–Thm2ex4r (5’-GTT CAA GGC CAC CTT TGC T-3’) and was expected to produce a WT band of 1,000bp. PCR amplicons with a lower molecular weight indicate evidence of NHEJ in exon 4. Primers designed to detect the large exon 4-8 deletion flank exon 4 and intron 8 include F-Thm2ex4b (5’-GGA GAG CAG CTT GAA GGA AA-3’) and R-Thm2ex4b (5’-GTC ACG GCT GGT GTG ATT C-3’). A PCR amplicon was expected of approximately 216bp, only in the event of NHEJ. To detect the presence of a WT allele, a separate PCR reaction was performed using primers within exon 6, F-WT (5-AAC TTC CTG CCC GCT TTA GT-3’) and R-WT (5’-GTG TCA GAT ACC CTG GAA CCA GAG-3’). In the presence of a WT allele, this PCR reaction yields an amplicon of approximately 461bp.

### Sequencing of Thm2 knockout alleles

PCR products were run on an agarose gel, excised, and extracted using the Qiaex II DNA extraction kit (Qiagen, 20021). Samples were sequenced by Genewiz. Sequencing results were analyzed to determine the presence or absence of stop codons resulting from NHEJ events. A line harboring a deletion from exon 4 to intron 8, producing a stop codon in exon 4 was chosen, with the allele designation of *Ttc21a*^∆*4-8*^. The founder of this line was mated to FVB mice to expand the line. This line was maintained on the C57BL6/J/FVB mixed genetic background.

### Analysis of mouse embryos and generation of mouse embryonic fibroblasts

Timed matings were performed between *Ttc21a*^∆*4-8/+*^; *Thm1*^*aln/+*^ mice. The *aln* allele of *Thm1* results in absence of protein, and thus acts like a null allele (11). Visualization of a vaginal plug was designated as embryonic day (E) 0.5. Mouse embryos were dissected at E10.5, E12.5 and E14.5 using a Leica dissection microscope. Tails of mouse embryos were collected for genomic DNA extraction and genotyping. To generate MEF, embryos were eviscerated. In a fresh 10cm cell culture plate, an individual carcass was minced with a razor in 0.25% trypsin-EDTA, then media (DMEM containing 10% FBS and penicillin streptomycin antibiotics) was added. Cells were grown to confluency, then trypsinized and plated for an experiment. Cells were passaged maximally up to two times.

### Cilia assembly and disassembly assays

Cells were plated on 12mm poly-L-lysine-coated coverslips in a 24-well plate with DMEM medium containing 10% FBS and pen/strep antibiotics. Two days after cells reached 100% confluency, medium was replaced with serum-free medium for 24 hours to induce ciliogenesis. Cells were fixed, immunostained for ciliary markers, imaged and quantified for presence of cilia.

To measure cilia disassembly, cells were plated on 12mm poly-lysine coated glass coverslips in a 24-well plate with complete medium containing 10% FBS. Two days after cells reached 100% confluence, media was replaced with serum-free media for 24 hours to bring cells to G0 and induce ciliogenesis. Following 24-hour serum starvation, cells were cultured in media containing 10% FBS for 2 hours to induce cilia disassembly (20). Cells were fixed, immunostained for ciliary markers, imaged and quantified for presence of cilia.

### Immunofluorescence

Cells were washed with PBS, then fixed with 4% paraformaldehyde/0.2% triton X-100 in PBS for 10 minutes at room temperature. Cells were washed again with PBS, and blocked with 1% BSA in PBS for 1 hour. Cells were then incubated with antibodies against SMO (Abcam), IFT52, IFT81, IFT88, IFT140, BBS2, BBS5 (Proteintech), ARL13B and INNP5E (Proteintech) and acetylated α-tubulin (Sigma) overnight at 4° C. Following 3 washes in PBS, cells were incubated with anti-rabbit AF488 and anti-mouse AF594 (InVitrogen Technologies) for 30 minutes at room temperature. Cells were washed 3X in PBS, and mounted with Vectashield containing 4,6-diamidino-2-phenylindole (DAPI) (Vector Laboratories). Immuno-labeled cells were viewed and imaged using a Leica TCS SPE confocal microscope configured on a DM550 Q upright microscope.

### qPCR

RNA was extracted using Trizol (Life Technologies), then reverse transcribed into cDNA using Quanta Biosciences qScript cDNA mix (VWR International). qPCR for *Gli1* was performed using Quanta Biosciences Perfecta qPCR Supermix (VWR International) in a BioRad CFX Connect Real-Time PCR Detection System. Primers used were m*Gli1* (Forward: 5’-CTGACTGTGCCCGAGAGTG-3’; Reverse: 5’-CGCTGCTGCAAGAGGACT-3’). qPCR was performed using RNA lysates from three lines for each genotype.

## Results

### *Thm2* interacts with *Thm1* in embryogenesis

Timed matings of *Thm2*^∆*4-8/+*^;*Thm1*^*aln/+*^ intercrosses were performed. At E10.5, E12.5 and E14.5, pregnant females were dissected and their embryos analyzed. *Thm2*-null (*Thm2*^∆*4-8/*∆*4-8*^) and *Thm2*-null;*Thm1+/−*(*Thm2*^∆*4-8/*∆*4-8*^;*Thm1*^*aln/+*^) embryos were normal, while *Thm1*-null (*Thm1*^*aln/aln*^) and *Thm1*; *Thm2* double knock-out (*Thm2*^∆*4-8*/∆*4-8*^;*Thm1*^*aln/aln*^) embryos showed polydactyly and occasional exencephaly. At E10.5 and E12.5, the number of *Thm1*-null mutants and *Thm1; Thm2* double knock-outs (dko) was consistent with a Mendelian inheritance pattern, but at E14.5, the frequency of *Thm1; Thm2* dko embryos was reduced (Table 1). *Thm1* deletion causes perinatal lethality (11). Thus, the reduced number of *Thm1; Thm2* double mutants at E14.5 suggests that additional loss of *Thm2* exacerbates the *Thm1*-null developmental phenotype causing mid-gestational lethality.

**Table 1.**
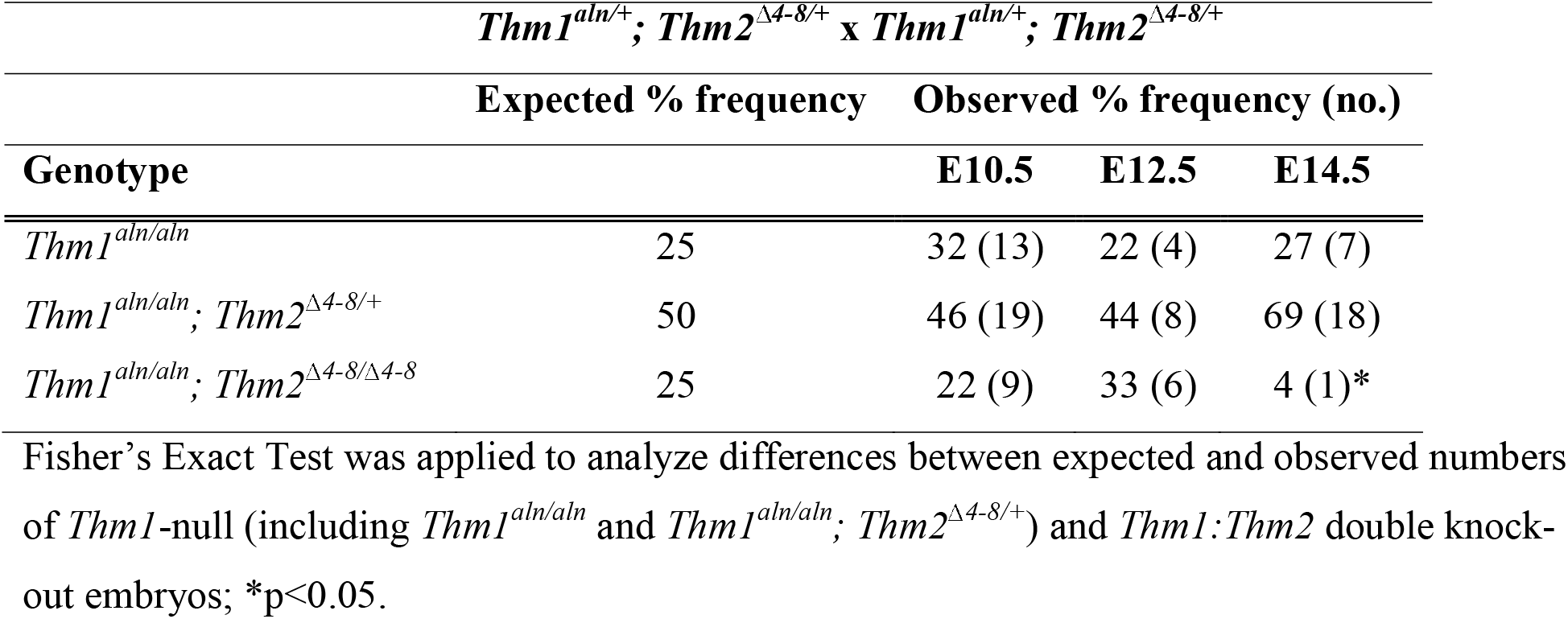
Frequency of live *Thm1;Thm2* double knock-out embryos

### *Thm1* regulates ciliogenesis

We derived mouse embryonic fibroblasts (MEF) from E12.5 *Thm2*-null, *Thm2*-null;*Thm1*^*+/−*^ (triple allele mutant), *Thm1*-null (ko), and *Thm1*; *Thm2* double-knockout (dko) mice. Following 24 hours of serum starvation, *Thm1* ko cells exhibited shorter ciliary length (mean length of 3.5µm) than control cells (*Thm2+/−*; mean length of 5.2µm), while the loss of *Thm2* did not alter cilia length (*Thm2*-null and *Thm2*-null;*Thm1+/−*mutants; mean lengths of 5.3µm and 5.2µm, respectively; Figures 1A and 1B).

**Figure 1.**
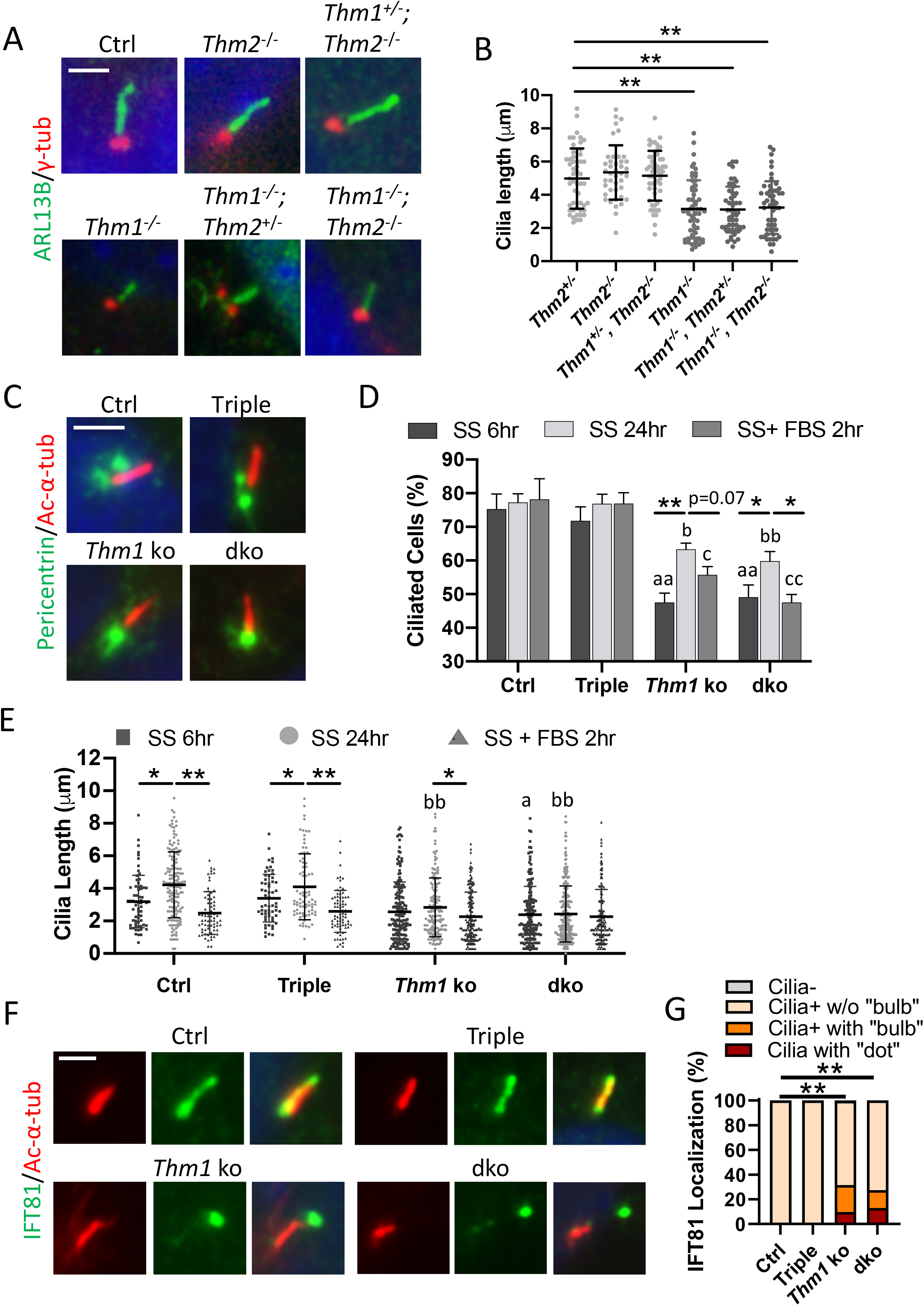
*Thm2* alters stability of pre-established cilia. (A) Immunostaining for ARL13B (green) and γ-tubulin (red). Scale bar = 5 µm. (B) Cilia length. 2-3 cell lines per genotype, and ≥40 cilia/genotype were randomly selected per experiment. Each dot represents an individual cilium length. Bars represent mean ± SD. (C) Immunostaining for acetylated α-tubulin (red) and pericentrin (green). Scale bar = 5 µm. (D) Percentage of ciliated MEF and (E) cilia length following 6hr serum starvation, 24hr serum starvation, and 24hr serum starvation +2hr FBS. Five-to-ten fields were analyzed per condition per experiment, and included ≥ 65 ciliated cells per condition per genotype. Bars represent mean ± SD from 2-3 independent experiments. Statistical significance was determined by ANOVA followed by Tukey’s or Dunnett’s test. *p<0.05; ** p<0.0001; ^a^ p<0.05; ^aa^ p<0.0001 compared to Ctrl MEF - 6hr serum starvation; ^b^ p<0.05; ^bb^ p<0.0001 compared to Ctrl MEF - 24hr serum starvation; ^c^ p<0.05; ^cc^ p<0.0001 compared to Ctrl MEF - 24hr serum starvation + 2hr FBS. (F) Immunostaining for IFT81 (green) and acetylated α-tubulin (red) following 24hr serum starvation + 2hr FBS. (G) Quantification of ciliary localization of IFT81, categorized as absent from cilia (Cilia-), present in cilia without a bulbous distal tip (Cilia + w/o bulb), and present in cilia with a bulbous distal tip (Cilia+ with bulb). Stacked bar graphs represent percentage of these occurrences. Total number of Ctrl, Triple, *Thm1* ko and dko cells quantified were 202, 205, 340 and 131, respectively, from 2-3 independent experiments. Statistical significance was determined by χ^2^ test. **p<0.0001

We next assessed capacity of mutant cells to undergo ciliogenesis. Loss of *Thm2* alone does not affect embryogenesis nor result in a phenotype at weanling age, but loss of *Thm2* together with one allele of *Thm1* (*Thm2*-null;*Thm1+/−*) causes a postnatal skeletal phenotype (data will be communicated in a separate manuscript). Therefore, all subsequent experiments were carried out using MEF of *Thm2*-null;*Thm1+/−*(triple allele) mutants rather than of *Thm2*-null cells. At 6 and 24 hours of serum starvation, percentage of ciliated cells and cilia length was measured in control (*Thm2*^∆*4-8/+*^), triple allele mutant (*Thm2*^∆*4-8*/∆*4-8*^;*Thm1*^*aln/+*^), *Thm1* ko(*Thm1*^*aln/aln*^) and *Thm1*; *Thm2* dko (*Thm1*^*aln/aln*^; *Thm2*^∆*4-8*/∆*4-8*^) MEF at 100% confluency. At 6 hours of serum starvation, approximately 75% of control MEF were ciliated. At 24 hours of serum starvation, percentage of ciliated control cells was not altered, but cilia length was increased (Figures 1C-1E). We posit that under our experimental conditions, early events in cilium formation, such as maturation and docking of the basal body, were completed at 6 hours of serum starvation, and that axoneme elongation occurred from 6 to 24 hours of serum starvation. In triple allele mutant MEF, percentage of ciliated cells and cilia lengths were similar to control MEF. In contrast, percentage of ciliated *Thm1*-null MEF was lower than control MEF (48% vs. 75%, respectively) at 6 hours serum starvation, and increased (to 64%) at 24 hours of serum starvation, suggesting absence of *Thm1* reduces and delays ciliogenesis. From 6 to 24 hours serum starvation, cilia length did not change in *Thm1*-null MEF, suggesting that events required for axoneme elongation are hindered by *Thm1* loss. A similar percentage of ciliated cells and cilia length was observed in *Thm1;Thm2* dko MEF as in *Thm1*-null MEF, suggesting that *Thm2* does not participate in cilia assembly.

### *Thm2* interacts with *Thm1* to regulate cilia disassembly

Primary cilia undergo cycles of assembly and disassembly in coordination with the cell cycle, and the cilia assembly:disassembly ratio regulates cilia length (21, 22). To examine cilia disassembly, serum was added back to the media for 2 hours following 24 hours of serum starvation. In control and triple allele mutant MEF, percentage of ciliated cells was not significantly altered (approximately 78%), but cilia length was decreased (Figures 1D and 1E). In *Thm1*-null MEF, the 2-hour serum addition resulted in a lower mean of percent ciliated cells (63% vs. 55%) although this reduction did not reach statistical significance. Additionally, cilia length was not altered. In *Thm1; Thm*2 dko MEF, serum addition significantly reduced percent ciliated cells, but did not modify cilia length. These data suggest that the loss of *Thm1* and *Thm2* promotes cilia disassembly.

Following 2 hours of serum addition, cells were also immunostained for IFT-B protein, IFT81. This revealed the presence of IFT81 to be separate and distal to its axonemal localization (Figure 1F; cilia + dot). This observation may reflect the release of ciliary vesicles from the distal tip, a phenomenon that is termed decapitation or ectocytosis, and contributes to cilia disassembly (23). In *Thm1*-null and *Thm1*; *Thm2* dko cells, frequency of IFT81 localization that was separate from the axoneme was increased (Figure 1G), suggesting *Thm1* regulates this phenomenon.

### *Thm2* interacts with *Thm1* to regulate retrograde transport of IFT-B component, IFT81

We next investigated the roles of *Thm1* and *Thm2* in ciliary protein transport. Following 24 hours of serum starvation to maximize cilia length, cells were immunostained for IFT-B subunits, IFT81 and IFT52, and for anterograde IFT motor, KIF3A. Ciliary localization of IFT-B proteins was similar between control and triple allele mutant MEF, while *Thm1*-null and *Thm1;Thm2* dko MEFs exhibited aberrant accumulation of proteins in a bulbous distal tip, indicative of defective retrograde IFT (Figures 2A-2C). As described previously for IFT-A mutant cells (7), protein localization was classified as either absent in cilia (Cilia-), present in cilia without a bulb (Cilia w/o bulb), or present in cilia with a bulb (Cilia with bulb). Across all genotypes, virtually 100% of cilia showed the presence of IFT81, IFT52 and KIF3A. However, while most control and triple allele mutant MEF showed localization of IFT81, IFT52 and KIF3A in cilia without a bulb, approximately 50% of *Thm1*-null and *Thm1;Thm2* dko MEF showed localization of IFT81, IFT52 and KIF3A in a bulbous ciliary distal tip. Additionally, *Thm1*; *Thm2* dko MEF displayed a higher percentage of cells with IFT81 localizing to a ciliary bulbous distal tip than *Thm1*-null MEF (52% vs. 43%; Figure 2A). These data indicate that THM1 is required for retrograde transport of these IFT-B proteins, and that THM2 enhances THM1-mediated retrograde transport of IFT81.

**Figure 2.**
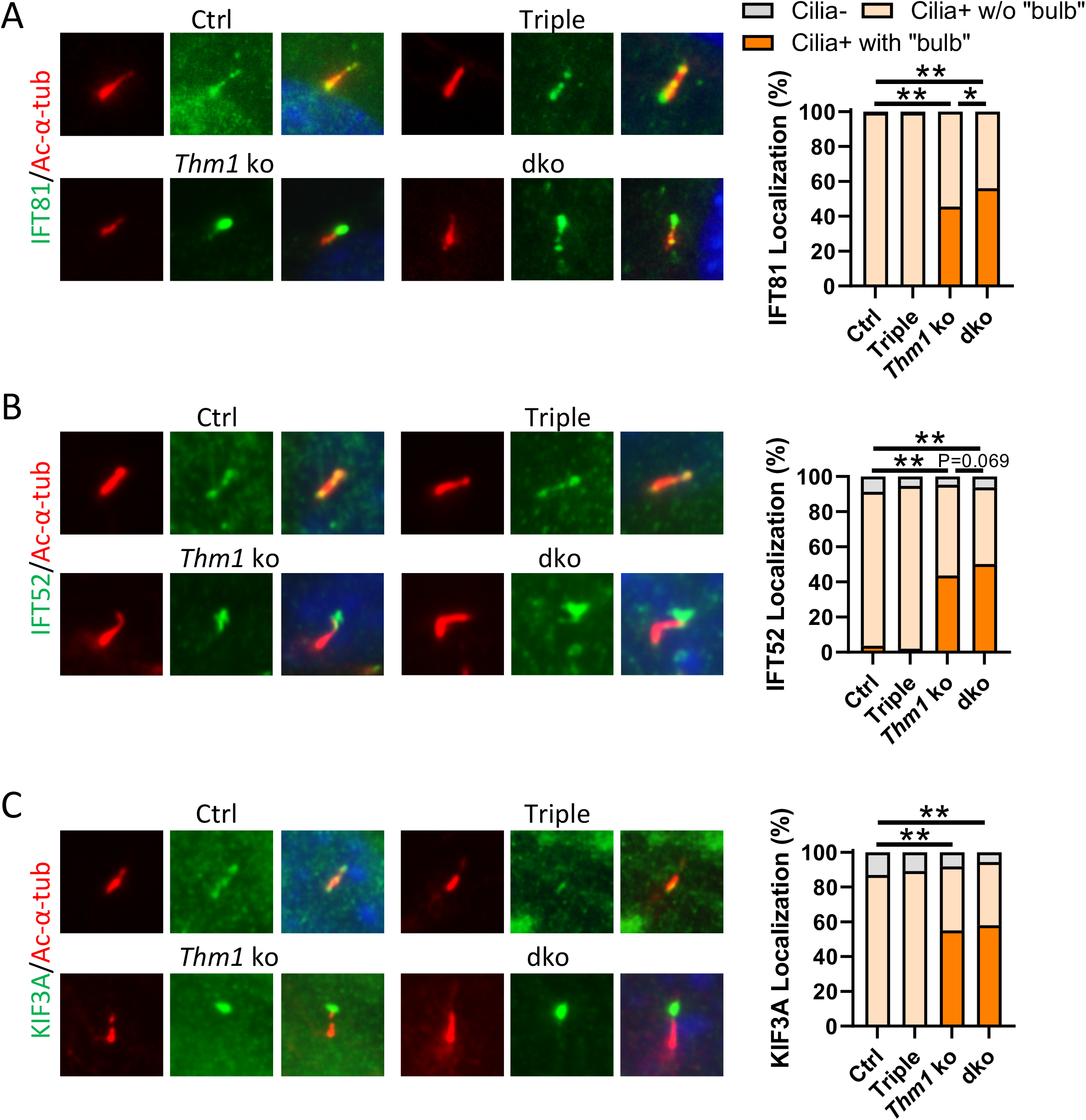
*Thm2* interacts with *Thm1* to regulate retrograde transport of IFT-B complex. Immunostaining and quantification for (A) IFT81 (green); (B) IFT52 (green) and (C) KIF3A (green). Ciliary localization of IFT81, IFT52 and KIF3A was categorized as absent from cilia (Cilia-), present in cilia without a bulbous distal tip (Cilia + w/o bulb), and present in cilia with a bulbous distal tip (Cilia+ with bulb). Stacked bar graphs represent percentage of these occurrences. Total number of Ctrl, Triple, *Thm1* ko and dko cells quantified were 150, 151, 200 and 200, respectively, were IFT81; 220, 244, 407 and 408, respectively, for IFT52; were 220, 244, 407 and 408, respectively, for KIF3A, from 2-3 independent experiments. Statistical significance was determined by χ^2^ test. *p<0.05; **p<0.0001

### *Thm1* regulates ciliary entry and retrograde transport of IFT-A component, IFT140

We next examined ciliary localization of IFT-A component, IFT140. Approximately 90% of cilia of control and triple allele mutant MEF showed the presence of IFT140, while only 65% of *Thm1*-null and *Thm1*; *Thm2* dko MEF had IFT140-positive cilia. Additionally, approximately 35% of *Thm1*-null and *Thm1*; *Thm2* dko cilia showed IFT140 localization in a bulbous distal tip (Figures 3A and 3B). These data indicate that THM1 mediates both ciliary entry and retrograde transport of an IFT-A protein.

**Figure 3.**
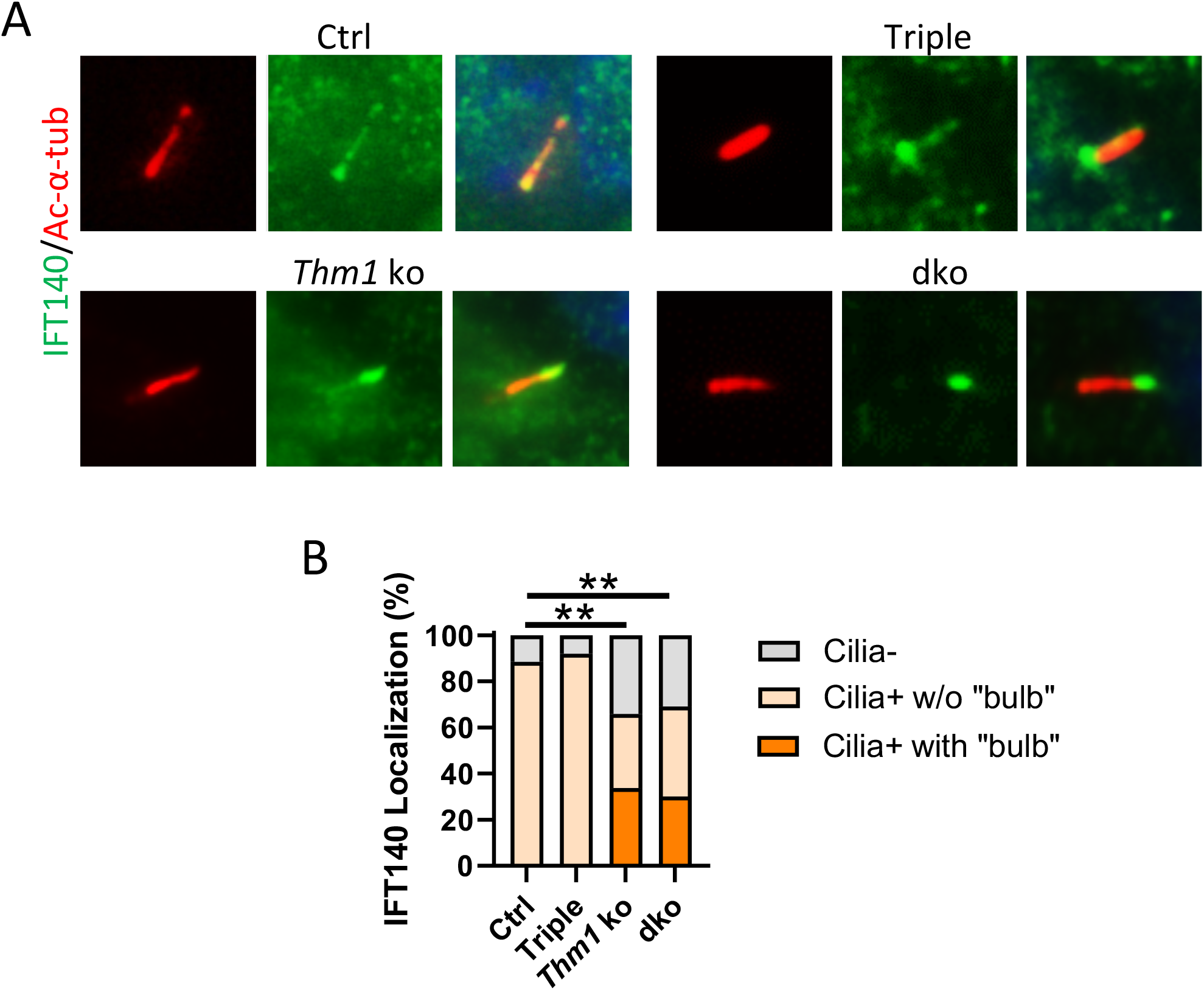
*Thm1* regulates retrograde transport and ciliary entry of IFT-A component, IFT140. (A) Immunostaining for IFT140 (green) and acetylated-α-tubulin (red). (B) Quantification of IFT140 ciliary localization. Total number of Ctrl, Triple, *Thm1* ko and dko cells quantified were 224, 274, 386 and 385, respectively, from 2-3 independent experiments. Statistical significance was determined by χ^2^ test. **p<0.0001

### *Thm1* mediates retrograde transport of BBSome subunits

The BBSome is an 8-subunit complex, which acts like an adaptor between IFT complexes and protein cargo within cilia. BBSome subunits are normally rapidly exported from cilia (7, 24, 25). Consistent with this, we observed light ciliary staining of BBS2 in approximately 80% of control and triple allele mutant MEF (Figures 4A-4C) and light ciliary staining of BBS5 in approximately 62% of control and triple allele mutant MEF (Figures 4D-4F). However, approximately 95% of *Thm1*-null and *Thm1*; *Thm2* dko MEF showed ciliary presence of BBS2 and BBS5, and approximately 20% showed BBS2 and BBS5 localization in bulb-like structures at the distal tip (Figures 4B and 4E). *Thm1*-null and *Thm1*; *Thm2* dko cilia also showed increased intensity of BBS2 and BBS5 immunofluorescence relative to control and triple allele mutant cilia (Figures 4C and 4F). These data indicate that THM1 loss impairs retrograde IFT of the BBSome.

**Figure 4.**
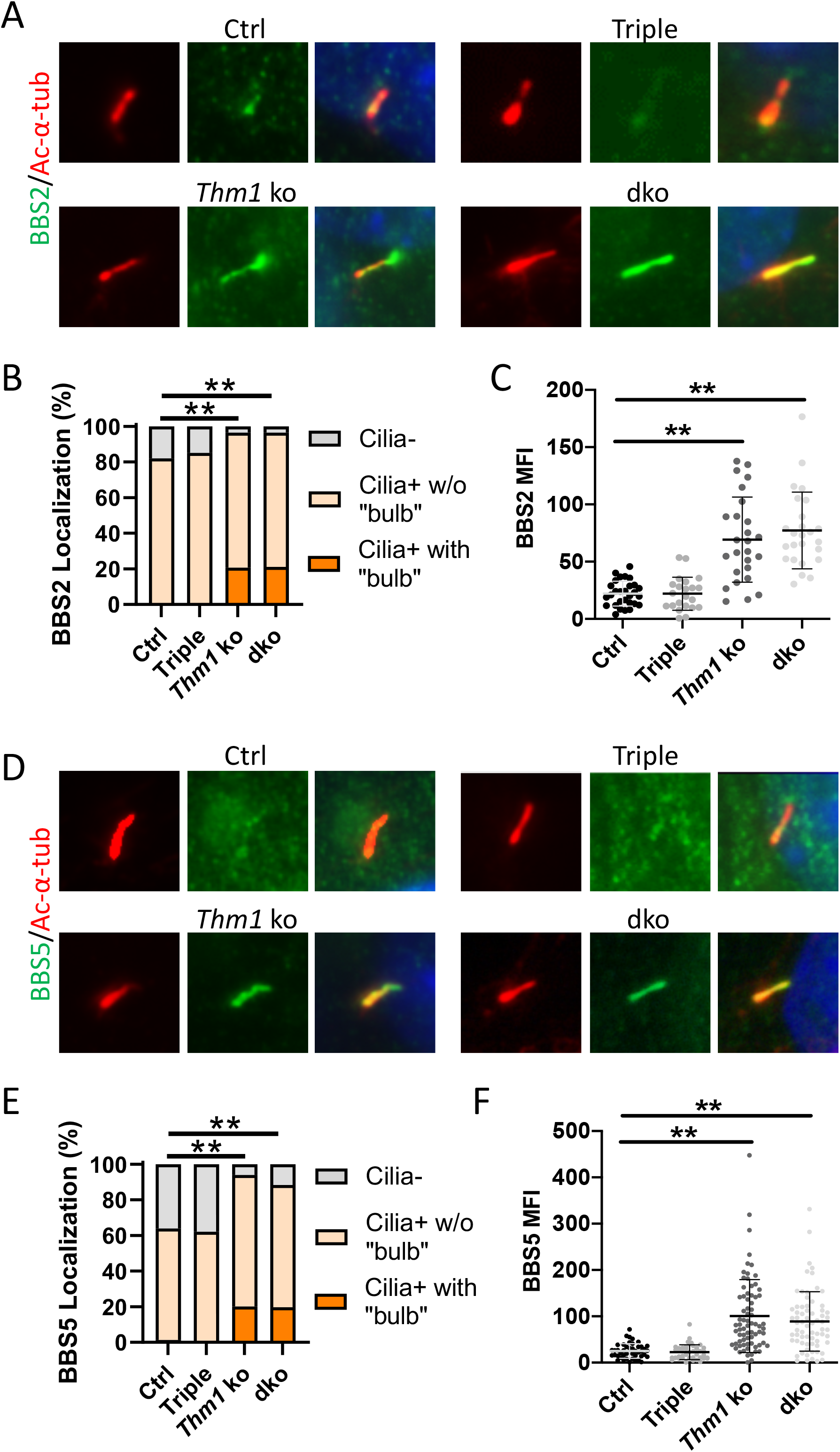
BBSome subunits are increased in *Thm1*-null cilia. (A) Immunostaining for BBS2 (green) and ac-α-tubulin (red). (B) Quantification of BBS2 ciliary localization. Total number of Ctrl, Triple, *Thm1* ko and dko cells quantified were 183, 167, 334 and 223, respectively, from 2-3 independent experiments. Statistical significance was determined by χ^2^ test. (C) Fluorescence intensity of BBS2 in cilia. Each dot represents an individual cilium. Bars represent mean ± SD. Statistical significance was determined by ANOVA followed by Tukey’s test. (D) Immunostaining for BBS5 (green) and ac-α-tubulin (red). (E) Quantification of BBS5 ciliary localization. Total number of Ctrl, Triple, *Thm1* ko and dko cells quantified were 219, 261, 229 and 217, respectively, from 3 independent experiments. Statistical significance was determined by χ^2^ test. (E) Fluorescence intensity of BBS5 in cilia. Each dot represents an individual cilium. Bars represent mean ± SD. Statistical significance was determined by ANOVA followed by Tukey’s test. **p<0.0001

### *Thm2* interacts with *Thm1* to mediate ciliary localization of INPP5E

Since the IFT-A complex mediates ciliary entry of membrane-associated proteins (6–8), we next examined ciliary localization of ARL13B and INPP5E. While 100% of control and triple allele mutant MEF showed ciliary presence of ARL13B, approximately 90% of *Thm1*-null and *Thm1*; *Thm2* dko cilia were positive for ARL13B (Figures 5A and 5B). Additionally, immunofluorescence intensity was decreased by almost 50% in *Thm1*-null and *Thm1*; *Thm2* dko cilia relative to control and triple allele mutant cilia (Figure 5C). Further, while 100% of control and triple allele mutant MEF showed ciliary presence of INPP5E, only 50% of *Thm1*-null MEF and 40% of *Thm1*; *Thm2* dko MEF had INPP5E-positive cilia (Figures 5D and 5E). These data suggest *Thm1* regulates ciliary entry of ARL13B and INPP5E, and that *Thm2* enhances *Thm1*-mediated ciliary entry of INPP5E.

**Figure 5.**
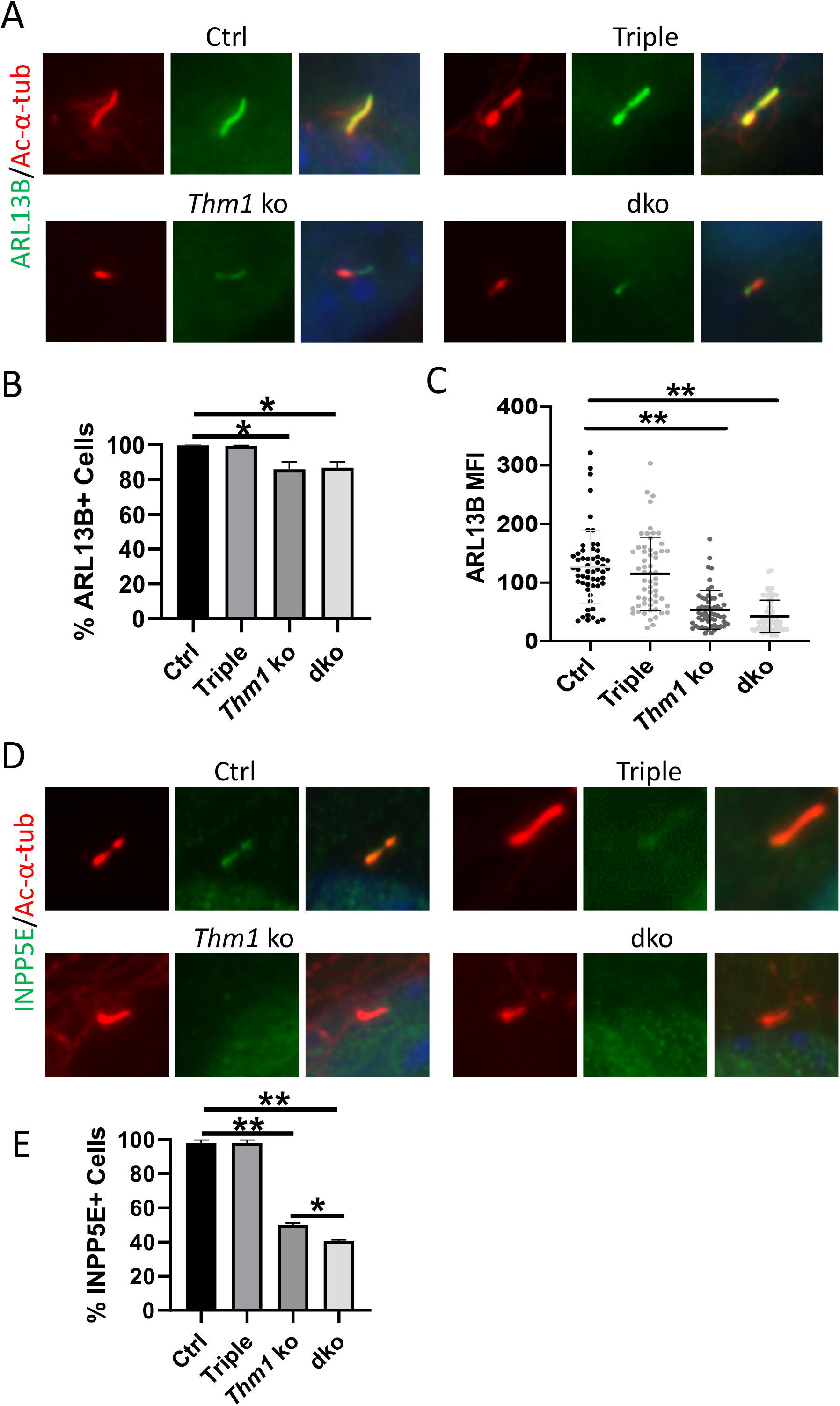
*Thm2* interacts with *Thm1* to mediate ciliary localization of ARL13B and INPP5E. (A) Immunostaining for ARL13B (green) and ac-α-tubulin (red). (B) Quantification of ARL13B positive cilia. ≥ 150 cells/genotype from 3 independent experiments were quantified. (C) Fluorescence intensity of Arl13b in cilia. Each dot represents an individual cilium. Bars represent mean ± SD. (D) Immunostaining for INPP5E (green) and ac-α-tubulin (red). (E) Quantification of INPP5E positive cilia. ≥ 150 cells/genotype from 3 independent experiments were quantified. Statistical significance was determined by ANOVA followed by Tukey’s test. *p<0.05; **p<0.0001

### *Thm2* interacts with *Thm1* to regulate Hh signaling

During activation of the mammalian Hedgehog (Hh) signaling pathway, the Smoothened (SMO) signal transducer enriches in the primary cilium (26). In control MEF, treatment with SMO agonist, SAG, resulted in presence of SMO in 60% of cilia, compared to 20% of cilia in untreated MEF (Figures 6A and 6B). Treatment with SAG also induced expression of *Gli1*, a transcriptional target and reporter of Hh activity (27). In triple allele mutant MEF, SAG treatment resulted in a ciliary enrichment of SMO similar to control cells (Figure 6B), yet *Gli1* expression was reduced relative to control MEF (Figure 6C). In untreated *Thm1*-null and *Thm1*; *Thm2* dko MEF, SMO was present in 90% of cilia, and 20% showed SMO accumulation in a bulbous distal tip (Figure 6A). With SAG treatment, approximately 90% of cilia were SMO+, unchanged from untreated cells, but approximately 30% of cilia had SMO localized in a bulbous distal tip (Figure 6B). SAG-induced *Gli1* expression in *Thm1*-null and *Thm1*; *Thm2* dko MEF was minimal to almost absent (Figure 6C). Thus, *Thm1* regulates retrograde transport of SMO, and also Hh pathway activity downstream of SMO ciliary localization. Further, *Thm2* interacts with *Thm1*, to positively regulate Hh signaling downstream of SMO ciliary localization.

**Figure 6.**
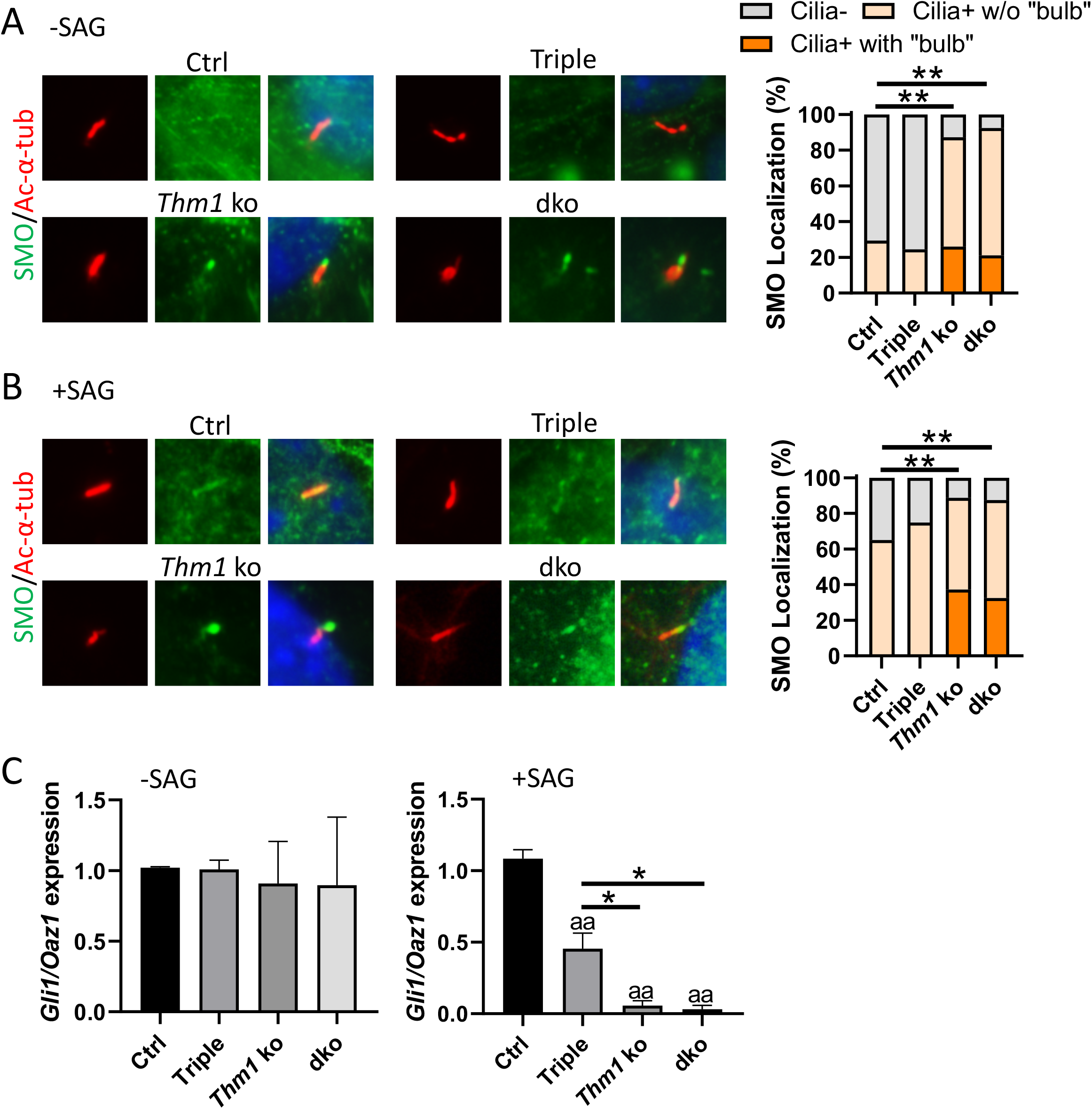
*Thm2* interacts with *Thm1* to regulate Hh signaling. Immunostaining for SMO and acetylated-α-tubulin of serum-starved MEF treated with (A) DMSO or (B) SAG for 8 hr, with quantification of SMO ciliary localization. Total number of Ctrl, Triple, *Thm1* ko and dko cells quantified in (A) were 191,135, 235 and 223, respectively, and in (B) were 203, 242, 363 and 200, respectively, from 2-3 independent experiments. Statistical significance was determined by χ^2^ test. (C) qPCR for *Gli1* on RNA extracts of MEF cultured in 1% FBS overnight, then treated with DMSO or SAG for 48 h in 1% FBS medium. *Gli1/Oaz1* transcript ratios of control cells were set to 1. Graphs represent mean ± SEM. Three MEF lines per genotype were used. Statistical significance was determined by ANOVA followed by Tukey’s test. *p<0.05; **p<0.0001; ^aa^ p<0.0001 compared to Ctrl

## Discussion

In this study, we reveal that *Thm1* loss decreases cilia assembly, and that loss of both *Thm1* and *Thm2* increases cilia disassembly. Since the ratio of cilia assembly:disassembly governs cilia length, this may account for the shorter cilia length in *Thm1*-null and *Thm1;Thm2* dko MEF. Further, in a *Thm1*-dependent manner, *Thm2* also regulates retrograde IFT, and Hh signaling activity downstream of SMO ciliary translocation.

To our knowledge, this is the first time that a role for IFT-A in cilia disassembly is shown. In control MEF, two hours of serum restimulation following 24hrs of serum starvation did not alter percentage of ciliated cells, but decreased cilia length. Conversely, in *Thm1; Thm*2 dko cells, two hours of serum restimulation decreased percentage of ciliated cells, while cilia length was unchanged. This difference between control and mutant cells may indicate that in the presence of dysfunctional IFT-A, the mechanism of cilia disassembly is shifted. Several mechanisms contribute to cilia disassembly. These include cilia resorption, which gradually shortens cilia length; complete cilia shedding, which cleaves the entire cilium; and a combination of both mechanisms (28). The reduction in cilia length in control MEF suggests gradual cilia resorption may be a predominant mechanism within 2hrs of FBS stimulation, whereas the unaltered cilia length together with less ciliated *Thm1; Thm*2 dko MEF may suggest whole cilium shedding may be more common when IFT-A is deficient. Additionally, we observed increased frequency of IFT81 localization that was separate and distal to the ciliary axoneme in *Thm1*-null and *Thm1; Thm*2 dko cells after 2 hour of serum addition. This IFT81 localization may present decapitation or ectocytosis (23). The reduced ciliary INPP5E in *Thm1*-mutant and *Thm1; Thm*2 dko MEF could promote ectocytosis, since *Inpp5e*-null MEF have enhanced ectocytosis (23). Ectocytosis precedes cilia resorption (23), thus cilia resorption may also occur in *Thm1*-null and *Thm1; Thm*2 dko cells, but may be more evident beyond 2 hours of serum restimulation. Further studies are required to determine the mechanisms by which IFT-A dysfunction regulates cilia disassembly.

The IFT-A complex can be subdivided into core and peripheral subcomplexes, comprised of IFT122/IFT140/IFT144 and of IFT42/IFT121/IFT139, respectively (18). Depletion of core versus peripheral subcomplex components can result in different phenotypes. For instance, *Ift144*-null RPE cells showed absence of SMO from cilia, suggesting defective ciliary import, while *Ift139-*null RPE cells accumulated SMO in cilia, indicating defective retrograde IFT (18). These contrasts may reflect differences in cargos carried by the subcomplexes or by the individual IFT-A proteins. As a paralog of *Thm1*, *Thm2* is likely a component of the IFT-A peripheral subcomplex. Consistent with this, *Thm1*-null and *Thm1;Thm2* dko MEF have ciliary protein trafficking defects similar to those of RPE cells depleted of another peripheral subcomplex component, *Ift121/WDR35* (7). This includes defective retrograde transport of IFT and BBSome proteins and impaired ciliary import of IFT-A protein, IFT140, and of membrane proteins, ARL13B and INPP5E. Similar also to IFT121/WDR35-depleted cells, *Thm1*-null MEF showed both reduced and delayed ciliogenesis. Reduced ciliary ARL13B in *Thm1*-null MEF may contribute to the reduced cilia assembly, since ARL13B is essential for ciliary membrane extension that is coupled to axoneme elongation (29).

Although ciliary presence of SMO was increased in *Thm1*-null and *Thm1*; *Thm2* dko MEF, *Gli1* expression was not induced by SAG treatment (Fig 6C), suggesting THM1 is required for Hh pathway activation downstream of SMO ciliary localization. This result was unexpected since we previously observed increased *Gli1* and *Ptch1* expression in E10.5 *Thm1*-null (*Thm1*^*aln/aln*^) whole-mount mouse embryos, and ventralization of the neural tube of E9.5 and E10.5 *Thm1*-null embryos indicating enhanced activation of the Hh pathway (11). Two possibilities may explain this discrepancy. Additional signals or cell-cell interactions may be present *in vivo*, resulting in enhanced activation of the pathway during development. Alternatively, a study has demonstrated that *Thm1* can act as either a positive or negative regulator of Hh signaling in a cell-specific manner, since deletion of *Thm1* resulted in reduced Hh signaling in glial cells during cerebellum development (30). We observed that SAG-treated triple allele mutant MEF had reduced pathway activation relative to control cells, indicating that in a *Thm1*-dose dependent manner, *Thm2* positively regulates Hh signaling downstream of SMO ciliary localization in MEF. In this light, absence of Hh activation in *Thm1*-null cells is consistent with *Thm1* acting as a positive modulator of Hh signaling in MEF. The factors that determine whether *Thm1* positively or negatively regulates the Hh pathway are yet to be defined.

**Figure 7.**
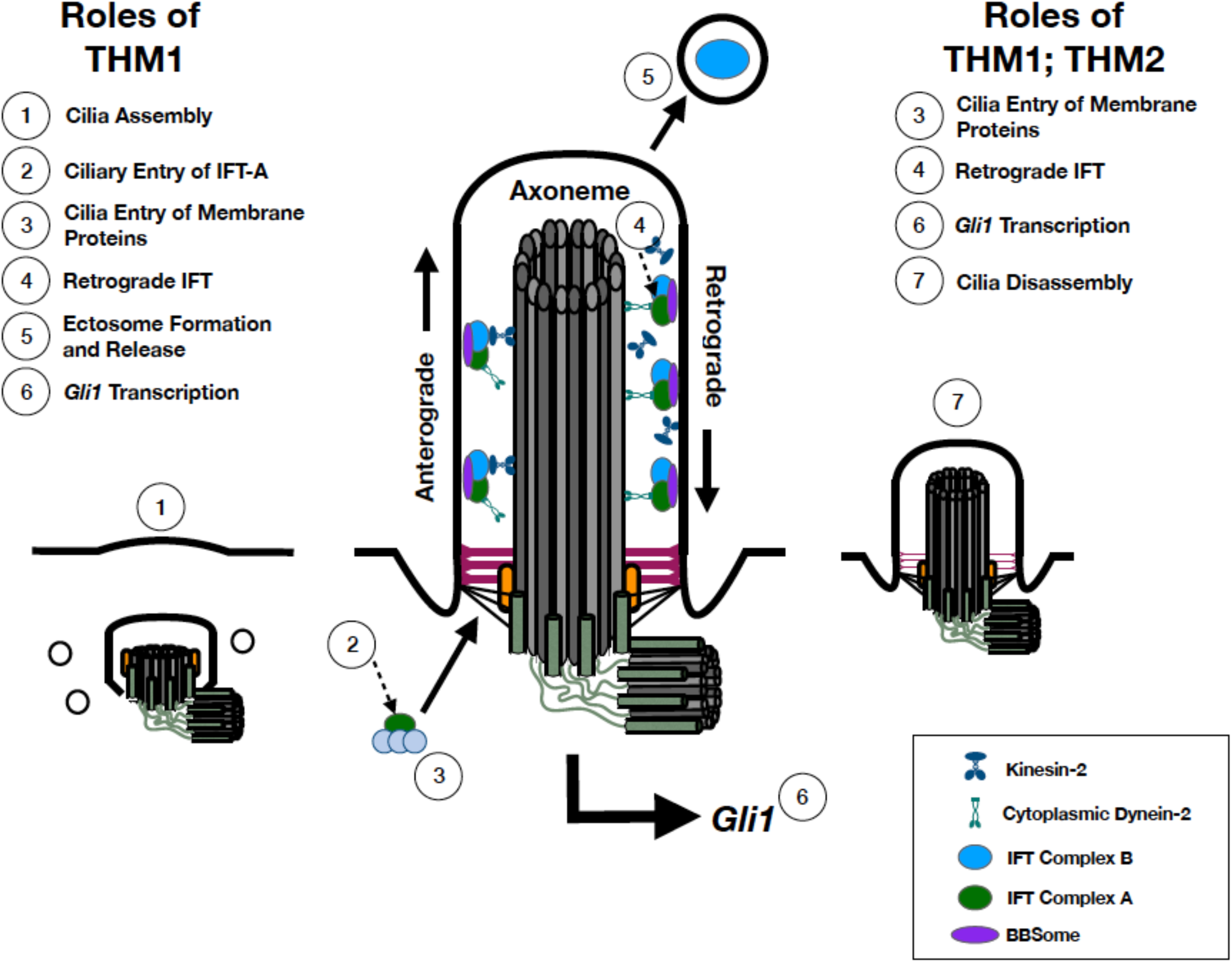
Ciliary roles of *Thm1* and *Thm2*. Model for *Thm1*, alone and together with *Thm2*, in regulating ciliogenesis, cilia disassembly, ciliary protein trafficking and Hh signaling in MEF.

Pathogenic mutations of *THM1* have been identified in approximately 5% of patients with ciliopathies (16). Interestingly, one-third of patients with a homozygous mutation in another ciliary gene also harbored a heterozygote *Thm1* mutation, suggesting that *THM1* mutations can have causal or modifying effects in human ciliopathies. More recently, mutations in *THM2* were reported in male individuals with subfertility (19). Our studies reveal that *Thm2* interacts with *Thm1*, and thus identifying mutations in *THM2* and *THM1* in the same individual may warrant investigation. Our findings that *Thm1*, alone and/or together with *Thm2*, regulates ciliary protein trafficking, cilia assembly and disassembly, Hh signaling, and development, provide insight into potential mechanisms underlying IFT-A related ciliopathies.

## Non-standard abbreviations

ARL13B: ADP Ribosylation Factor Like GTPase 13B
BBS: Bardet Biedl Syndrome
E: embryonic day
Hh: Hedgehog
INPP5E: Inositol Polyphosphate-5-Phosphatase E
IFT: intraflagellar transport
MEF: mouse embryonic fibroblasts
SAG: Smoothened agonist
SMO: Smoothened
*Thm1*: TPR-containing Hedgehog modulator 1
TPR: tetratricopeptide repeat

## Acknowledgements

We thank the KUMC Transgenic and Gene Targeting Institution Facility for generation of the *Thm2* mutant mice and acknowledge support of this Facility (Intellectual and Developmental Disabilities Research Center NIH U54 HD090216; KU Cancer Center NIH P30 CA168524; COBRE NIH P30 GM122731). This work was also supported by the National Institutes of Health [P30DK106912 to BAA]; [R01DK103033 to PVT].

## Author Contributions

WW and BAA performed experiments. WW, BAA, JLV and PVT analyzed data. WW, JLV and PVT designed research. TSP generated model. WW and PVT wrote the paper.

## References

1. Plotnikova, O. V., Pugacheva, E. N., and Golemis, E. A. (2009) Primary cilia and the cell cycle. Methods in cell biology 94, 137–160

2. Goetz, S. C., Ocbina, P. J., and Anderson, K. V. (2009) The primary cilium as a Hedgehog signal transduction machine. Methods in cell biology 94, 199–222

3. Tobin, J. L., and Beales, P. L. (2009) The nonmotile ciliopathies. Genet Med 11, 386–402

4. Brown, J. M., and Witman, G. B. (2014) Cilia and Diseases. Bioscience 64, 1126–1137

5. Keeling, J., Tsiokas, L., and Maskey, D. (2016) Cellular Mechanisms of Ciliary Length Control. Cells 5

6. Mukhopadhyay, S., Wen, X., Chih, B., Nelson, C. D., Lane, W. S., Scales, S. J., and Jackson, P. K. (2010) TULP3 bridges the IFT-A complex and membrane phosphoinositides to promote trafficking of G protein-coupled receptors into primary cilia. Genes Dev. 24, 2180–2193

7. Fu, W., Wang, L., Kim, S., Li, J., and Dynlacht, B. D. (2016) Role for the IFT-A Complex in Selective Transport to the Primary Cilium. Cell reports 17, 1505–1517

8. Picariello, T., Brown, J. M., Hou, Y., Swank, G., Cochran, D. A., King, O. D., Lechtreck, K., Pazour, G. J., and Witman, G. B. (2019) A global analysis of IFT-A function reveals specialization for transport of membrane-associated proteins into cilia. J. Cell Sci. 132

9. Jin, H., White, S. R., Shida, T., Schulz, S., Aguiar, M., Gygi, S. P., Bazan, J. F., and Nachury, M. V. (2010) The conserved Bardet-Biedl syndrome proteins assemble a coat that traffics membrane proteins to cilia. Cell 141, 1208–1219

10. Zhang, Q., Nishimura, D., Seo, S., Vogel, T., Morgan, D. A., Searby, C., Bugge, K., Stone, E. M., Rahmouni, K., and Sheffield, V. C. (2011) Bardet-Biedl syndrome 3 (Bbs3) knockout mouse model reveals common BBS-associated phenotypes and Bbs3 unique phenotypes. Proc. Natl. Acad. Sci. U. S. A. 108, 20678–20683

11. Tran, P. V., Haycraft, C. J., Besschetnova, T. Y., Turbe-Doan, A., Stottmann, R. W., Herron, B. J., Chesebro, A. L., Qiu, H., Scherz, P. J., Shah, J. V., Yoder, B. K., and Beier, D. R. (2008) THM1 negatively modulates mouse sonic hedgehog signal transduction and affects retrograde intraflagellar transport in cilia. Nat. Genet. 40, 403–410

12. Iomini, C., Li, L., Esparza, J. M., and Dutcher, S. K. (2009) Retrograde intraflagellar transport mutants identify complex A proteins with multiple genetic interactions in Chlamydomonas reinhardtii. Genetics 183, 885–896

13. Herron, B. J., Lu, W., Rao, C., Liu, S., Peters, H., Bronson, R. T., Justice, M. J., McDonald, J. D., and Beier, D. R. (2002) Efficient generation and mapping of recessive developmental mutations using ENU mutagenesis. Nature genetics 30, 185–189

14. Tran, P. V., Talbott, G. C., Turbe-Doan, A., Jacobs, D. T., Schonfeld, M. P., Silva, L. M., Chatterjee, A., Prysak, M., Allard, B. A., and Beier, D. R. (2014) Downregulating hedgehog signaling reduces renal cystogenic potential of mouse models. J. Am. Soc. Nephrol. 25, 2201–2212

15. Jacobs, D. T., Silva, L. M., Allard, B. A., Schonfeld, M. P., Chatterjee, A., Talbott, G. C., Beier, D. R., and Tran, P. V. (2016) Dysfunction of intraflagellar transport-A causes hyperphagia-induced obesity and metabolic syndrome. Dis Model Mech 9, 789–798

16. Davis, E. E., Zhang, Q., Liu, Q., Diplas, B. H., Davey, L. M., Hartley, J., Stoetzel, C., Szymanska, K., Ramaswami, G., Logan, C. V., Muzny, D. M., Young, A. C., Wheeler, D. A., Cruz, P., Morgan, M., Lewis, L. R., Cherukuri, P., Maskeri, B., Hansen, N. F., Mullikin, J. C., Blakesley, R. W., Bouffard, G. G., Gyapay, G., Rieger, S., Tonshoff, B., Kern, I., Soliman, N. A., Neuhaus, T. J., Swoboda, K. J., Kayserili, H., Gallagher, T. E., Lewis, R. A., Bergmann, C., Otto, E. A., Saunier, S., Scambler, P. J., Beales, P. L., Gleeson, J. G., Maher, E. R., Attie-Bitach, T., Dollfus, H., Johnson, C. A., Green, E. D., Gibbs, R. A., Hildebrandt, F., Pierce, E. A., and Katsanis, N. (2011) TTC21B contributes both causal and modifying alleles across the ciliopathy spectrum. Nat. Genet. 43, 189–196

17. Hirano, T., Katoh, Y., and Nakayama, K. (2017) Intraflagellar transport-A complex mediates ciliary entry and retrograde trafficking of ciliary G protein-coupled receptors. Mol. Biol. Cell 28, 429–439

18. Zhu, B., Zhu, X., Wang, L., Liang, Y., Feng, Q., and Pan, J. (2017) Functional exploration of the IFT-A complex in intraflagellar transport and ciliogenesis. PLoS genetics 13, e1006627

19. Liu, W., He, X., Yang, S., Zouari, R., Wang, J., Wu, H., Kherraf, Z. E., Liu, C., Coutton, C., Zhao, R., Tang, D., Tang, S., Lv, M., Fang, Y., Li, W., Li, H., Zhao, J., Wang, X., Zhao, S., Zhang, J., Arnoult, C., Jin, L., Zhang, Z., Ray, P. F., Cao, Y., and Zhang, F. (2019) Bi-allelic Mutations in TTC21A Induce Asthenoteratospermia in Humans and Mice. Am J Hum Genet 104, 738–748

20. Pugacheva, E. N., Jablonski, S. A., Hartman, T. R., Henske, E. P., and Golemis, E. A. (2007) HEF1-dependent Aurora A activation induces disassembly of the primary cilium. Cell 129, 1351–1363

21. Mirvis, M., Stearns, T., and James Nelson, W. (2018) Cilium structure, assembly, and disassembly regulated by the cytoskeleton. The Biochemical journal 475, 2329–2353

22. Spalluto, C., Wilson, D. I., and Hearn, T. (2013) Evidence for reciliation of RPE1 cells in late G1 phase, and ciliary localisation of cyclin B1. FEBS open bio 3, 334–340

23. Phua, S. C., Chiba, S., Suzuki, M., Su, E., Roberson, E. C., Pusapati, G. V., Setou, M., Rohatgi, R., Reiter, J. F., Ikegami, K., and Inoue, T. (2017) Dynamic Remodeling of Membrane Composition Drives Cell Cycle through Primary Cilia Excision. Cell 168, 264–279 e215

24. Eguether, T., San Agustin, J. T., Keady, B. T., Jonassen, J. A., Liang, Y., Francis, R., Tobita, K., Johnson, C. A., Abdelhamed, Z. A., Lo, C. W., and Pazour, G. J. (2014) IFT27 links the BBSome to IFT for maintenance of the ciliary signaling compartment. Developmental cell 31, 279–290

25. Liew, G. M., Ye, F., Nager, A. R., Murphy, J. P., Lee, J. S., Aguiar, M., Breslow, D. K., Gygi, S. P., and Nachury, M. V. (2014) The intraflagellar transport protein IFT27 promotes BBSome exit from cilia through the GTPase ARL6/BBS3. Developmental cell 31, 265–278

26. Corbit, K. C., Aanstad, P., Singla, V., Norman, A. R., Stainier, D. Y., and Reiter, J. F. (2005) Vertebrate Smoothened functions at the primary cilium. Nature 437, 1018–1021

27. Bai, C. B., Auerbach, W., Lee, J. S., Stephen, D., and Joyner, A. L. (2002) Gli2, but not Gli1, is required for initial Shh signaling and ectopic activation of the Shh pathway. Development 129, 4753–4761

28. Mirvis, M., Siemers, K. A., Nelson, W. J., and Stearns, T. P. (2019) Primary cilium loss in mammalian cells occurs predominantly by whole-cilium shedding. PLoS biology 17, e3000381

29. Lu, H., Toh, M. T., Narasimhan, V., Thamilselvam, S. K., Choksi, S. P., and Roy, S. (2015) A function for the Joubert syndrome protein Arl13b in ciliary membrane extension and ciliary length regulation. Dev. Biol. 397, 225–236

30. Driver, A. M., Shumrick, C., and Stottmann, R. W. (2017) Ttc21b Is Required in Bergmann Glia for Proper Granule Cell Radial Migration. J Dev Biol 5

